## Supplementary material for "Parietal alpha frequency shapes own-body perception by modulating the temporal integration of bodily signals": supllementary results and images

**Participants inclusion test**

Participants who do not experience a clear reliable rubber hand illusion are not able to make ownership discrimination in the body ownership judgment task. Therefore, participants underwent first an inclusion test to ensure that they could experience the illusion and be able to perform body ownership judgments in the main experiment. In the inclusion test, the visuo-tactile stimulation to induce the rubber hand illusion was identical to the one used in synchronous condition of the main experiment. The robots applied six synchronous touches to the index finger of the rubber hand and to the participant’s hidden real index finger for 12 seconds. At the end of the stimulation, participants were asked to complete a 9-items questionnaire to assess the ownership sensed over the rubber hand (**Attachment 1**). Participants were asked to indicate the extent of their agreement or disagreement with six statements using a seven-point Likert scale, ranging from −3 (“I completely disagree”) to +3 (“I completely agree”), with a response of 0 indicating “neither agreed nor disagreed”. Three of the statements examined the perception of the illusion (Q1–Q3) and the remaining two statements were designed to control suggestibility and task compliance (Q4-Q9). Our inclusion criteria for a rubber hand illusion strong enough for participation in the main psychophysics experiment were as follows: (i) a mean score for the illusion statements (Q1, Q2, and Q3) of greater than 1 and (ii) a difference between the mean score for the illusion items and the mean score for the control items of greater than 1.

**Supplementary figures and table**

**Experiment 2: correlations between resting-state IAFs and perceptual performance**

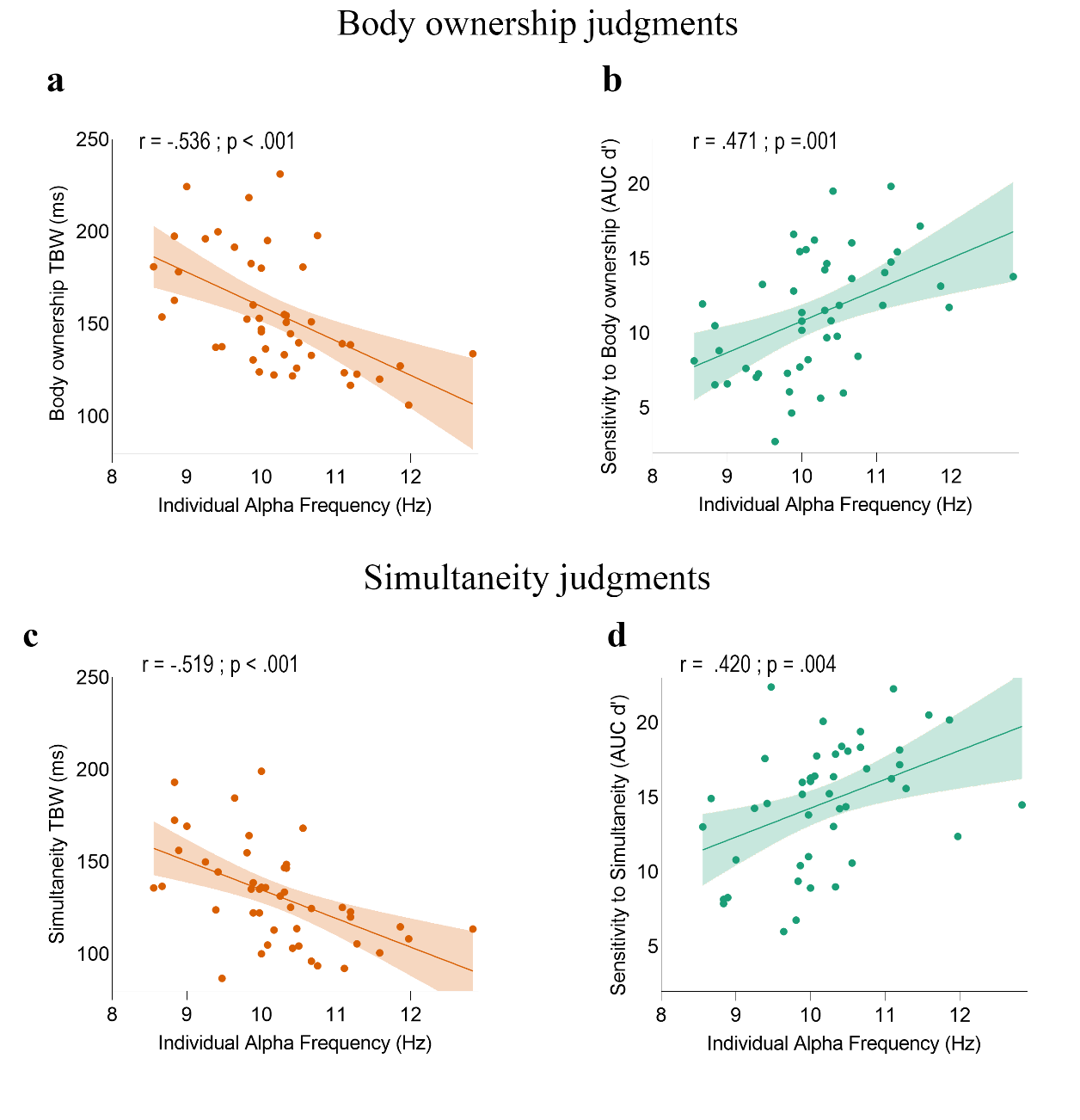

**Fig. S1.** **| Correlations between resting-state Individual Alpha Frequency (IAF) in the left parietal lobe and perceptual performance in Experiment 2. a, c,** Correlation between resting-state IAF measured in the left posterior parietal cortex and body ownership and simultaneity TBWs. **b, d,** Correlation between resting-state IAF in the left posterior parietal cortex and body ownership sensitivity and simultaneity sensitivity. The solid line represents the best-fitting regression. The shaded region reflects the 95% confidence interval.

**Experiment 3: results of the Bayesian computational model**

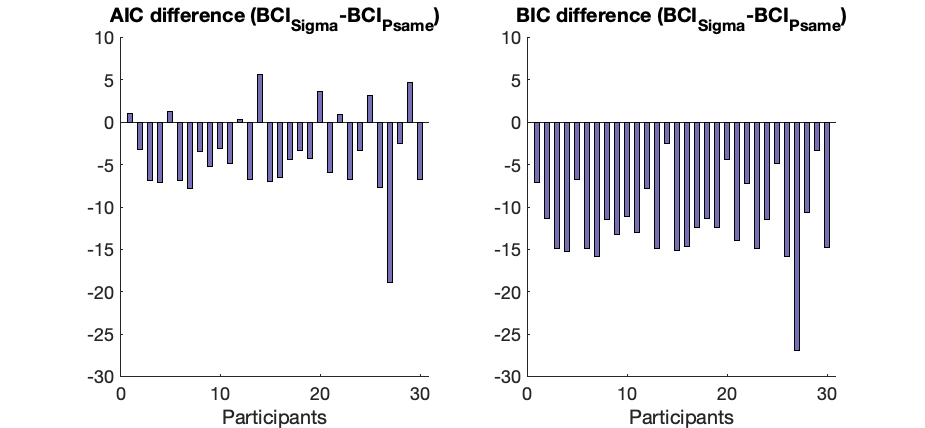

**Fig. S2 | Results of the computational modelling in Experiment 3**. **a,b**, Raw difference in AIC (**a**) and BIC (**b**) for each participant between the BCI_Sigma and the BCI_Psame models. A negative value means that the first model outperforms the latter.

**Experiment 3: Confidence intervals for the AIC and BIC differences between the two models**

**Table 1**: Bootstrapped confidence intervals (95% CI) for the AIC and BIC differences between a model that assumes a change in the level of sensory uncertainty and a model assuming a change in the a priori probability for a common source for the visual and tactile inputs. A negative value means that the first model is a better fit the data observed in the tACS Experiment

| Model comparison | AIC (95 % CI) | | | BIC (95 % CI) | | |
| --- | --- | --- | --- | --- | --- | --- |
| BCI_Sigma_-BCI_Psame_ | lower bound | raw sum | higher bound | lower bound | raw sum | higher bound |
|  | -165 | -112 | -61 | -407 | -355 | -304 |

**Supplementary results**

**Experiment 2: no correlation between alpha power and body ownership and simultaneity TBW**

We found no relationship between body ownership TBW and resting-state alpha power in either the left (Pearson’s r = -.004; p =.981; Spearman’s ρ = .114; p = .451) or right hemispheres (Pearson’s r = = -.026; p =.864; Spearman’s ρ = .066; p = .663). No significant correlation was found between body ownership TBW and task related alpha power measured during body ownership judgments (left parietal lobe: Pearson’s r = -.067, p = .659; Spearman’s ρ = -.005; p = .976; right parietal lobe: Pearson’s r = -.100, p = .507; Spearman’s ρ = -.039, p =.797).

Similarly, no relationship emerged between visuotactile simultaneity TBW and resting-state alpha power (left hemisphere: Pearson’s r = .131, p = .385; Spearman’s ρ = .170, p = .259; right hemisphere: Pearson’s r = .212, p = .157; Spearman’s ρ = .193, p = .199). No relationship emerged between simultaneity TBW and task-related alpha power measured during simultaneity judgments in either hemisphere (left: Pearson’s r = .129; p =.393; Spearman’s ρ = .089; p = .555; right: Pearson’s r = .173; p = .249; Spearman’s ρ = .201; p = .180).

**Experiment 2: no correlation between alpha power and body ownership and simultaneity sensitivities**

There was no correlation between sensitivity to body ownership and task-related alpha power, in either the left (Pearson’s r = -.083, p =.584; Spearman’s ρ = .022; p = .885) or right parietal lobes (Pearson r = .024, p =.874; Spearman’s ρ = .093; p = .537), nor was there any correlation between body ownership sensitivity and resting-state alpha power in the left (Pearson’s r = -.060, p = .692; Spearman’s ρ = -.055, p = .717) and right (Pearson’s r: -.024; p = .874; Spearman’s ρ = .011, p = .941) parietal lobe. Similarly, no correlation emerged between sensitivity to visuotactile simultaneity and task-related alpha power in either the left (Pearson’s r = -.185, p = .218; Spearman’s ρ = -.112, p = .457) or right (Pearson’s r = -.177, p = .238; Spearman’s ρ = -.147, p = .329) parietal cortex, nor with left (Pearson’s r = -.106, p = .482; Spearman’s ρ = -.104, p = .493) and right (Pearson’s r = -.223, p = .137; Spearman’s ρ = -.138, p = .359) resting-state alpha power.

**Experiment 2: no correlation between IAF and bias in body ownership and simultaneity judgments**

As a further control analysis, we also searched for possible associations between IAF and bias in body ownership and simultaneity judgments. The bias in body ownership does not correlate with task-related IAF measured during body ownership judgments, neither in the left (Pearson’s r = .041, p = .788; Spearman’s ρ = .030 p = .844) nor right parietal cortex (Pearson’s r = .046, p = .764; Spearman’s ρ = .052, p = .732). Similarly, no relationship emerged between the bias in body ownership judgments and parietal resting-state IAF (left: Pearson’s r = .121, p = .423; Spearman’ s ρ = .025, p =.867; right: Pearson’s r = -.006, p = .969; Spearman’ s ρ = -.029, p = .846). Moreover, no significant relationship emerged between bias in simultaneity judgments and parietal task-related IAF, either in the left (Pearson’s r = .119, p = .432; Spearman’s ρ = .153, p = .310) or right lobes (Pearson’s r = .129, p = .391; Spearman’s ρ = .109, p = .471). Similarly, no relationship emerged between bias in simultaneity task and resting-state IAF in the parietal left (Pearson’s r = -.116, p = .441; Spearman’ s ρ = -.075, p = .619) or right (Pearson’s r = .053, p = .727; Spearman’s ρ = -.023, p = .879) lobes. Therefore, our results suggest that IAF determines body ownership and simultaneity sensitivity, rather than being related to body ownership and simultaneity bias (e.g., more liberal or conservative responses).

**Experiment 2: correlation between body ownership and simultaneity TBWs and their perceptual sensitivities**

Experiment 2 successfully replicated the behavioural findings of Experiment 1. Body ownership and visuotactile simultaneity TBWs positively correlated (Pearson’s r = .563; p < .001; Spearman’s ρ = .590; p < .001; **Fig S3a**). Similarly, body ownership sensitivity correlated with simultaneity sensitivity (Pearson’s r = .652; p < .001; Spearman’s ρ = .662; p < .001; **Fig. S3b**). Using a paired t-test, we ascertain that body ownership TBW was significantly wider than that of simultaneity TBW (t = 5,912; p < .001). Finally, two ANOVAs conducted on body ownership and simultaneity sensitivities with asynchronies as factors revealed that both body ownership sensitivity (F _3,135_ = 586,953; p < .001; η_p_^2^ = .929) and simultaneity sensitivity (F _3,135_ = 529.727; p < .001; η_p_^2^ = .922) increased with the increasing of the asynchronies.

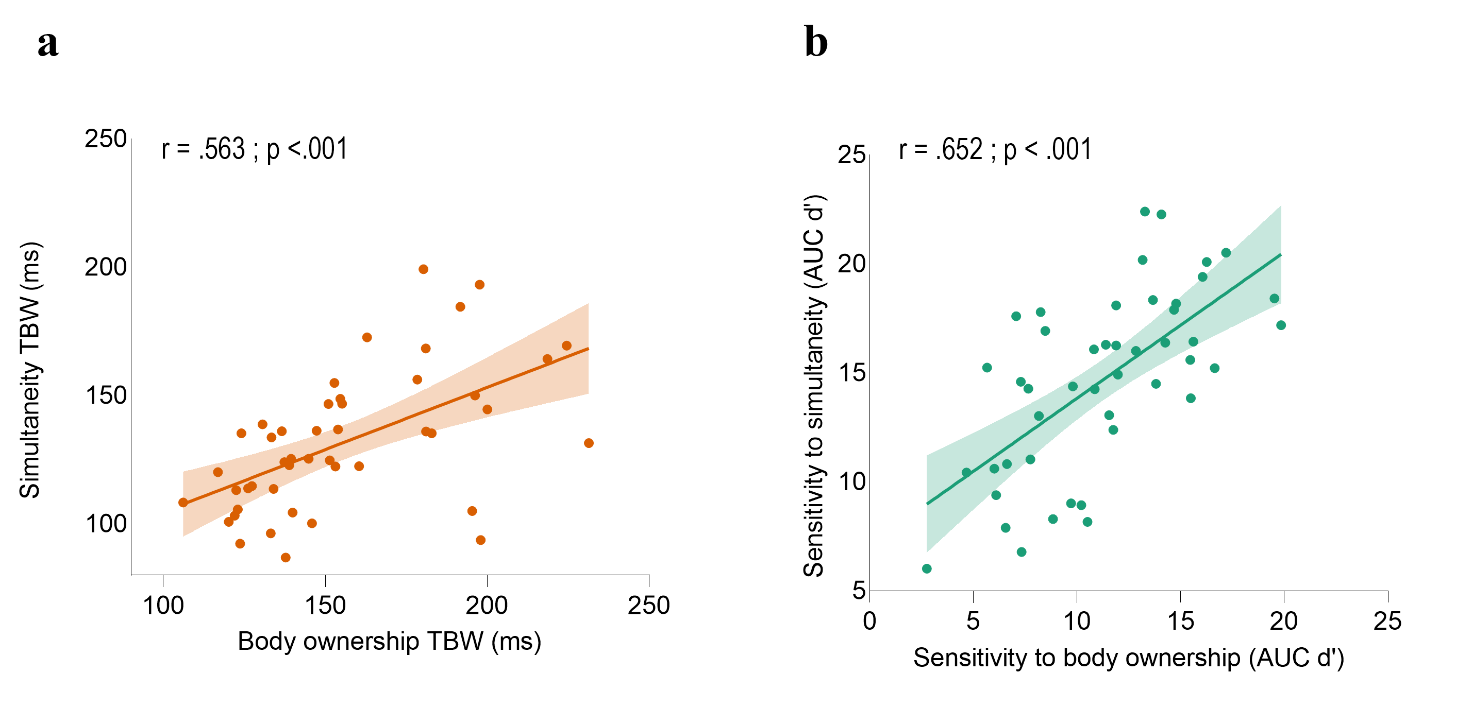
**Fig. S3 | Behavioural correlations in Experiment 2. a,** behavioural correlation between body ownership and visuo-tactile temporal binding windows (TBW). **b**, behavioural correlation between sensitivity to simultaneity and sensitivity to body ownership. In both graphs, the solid line represents the best-fitting regression. The shaded region reflects the 95% confidence interval. AUC denote the area under the curve of d’ scores measured for each asynchrony.

**Experiment 2: IAF modulation between the two tasks**

Next, we examined whether task-related changes in individual alpha frequency (IAF) reflect the different temporal demands of body ownership and simultaneity judgments. We hypothesized that alpha frequency would speed up during simultaneity judgments, as these require temporal segregation of visual and tactile input signals, i.e., higher temporal precision, to detect small delays. Previous studies have shown that alpha rates slow during tasks involving sensory integration and speed up when temporal segregation is required^1,2^. Therefore, we anticipate shorter alpha frequencies in the simultaneity task, reflecting the demand for precise timing in detecting simultaneity. In contrast, the body-ownership task lacks this temporal precision requirement and instead reflects a bodily illusion that builds over time through multisensory integration.

To this aim, we run an ANOVA with task (body ownership and simultaneity judgments) and laterality (left and right parietal lobes) as factors. We found that the IAF was slower when participants performed body ownership as compared to simultaneity judgments (F _1,45_ = 6,32; p = .016; η_p_^2^ = .123). Additionally, IAF was faster in the right as compared to left parietal lobe (F_1,45_ = 12,566; p = .001; η_p_^2^ = .218), with no significant interaction between hemisphere and task observed. In contrast, alpha power was not modulated by the task (F_1, 45_ = 0.241; p =.626; η_p_^2^ = 0.005), but it was lower in the left as compared to the right hemisphere (F_1, 45_ = 5.849; p = 0.020; η_p_^2^ = 0.115).

We reasoned that modulation of IAF between simultaneity and body ownership judgments could be associated with variations in TBW and sensitivity across the two tasks. To test this hypothesis, we calculated the difference between the TBW (Δ TBW) in body ownership and simultaneity judgments as a proxy for task-related TBW modulation, along with the difference in sensitivity (Δ d’) as a measure of sensitivity modulation. Similarly, we averaged the alpha frequency peak measured in the two parietal ROIs for each task. Then, the difference between the alpha peak measured in the body ownership and simultaneity judgments (Δ IAF) was taken as the measure of the degree to which each participant regulates the IAF in the two tasks. We found a positive correlation between Δ IAF and Δ TBW (Pearson’s r = .355; p = .016; Spearman’s ρ = .435, p = .003), as well as Δ IAF and Δ sensitivity (Pearson’s r = .306; p = .039; Spearman’ s ρ = .375, p = .010). Indeed, a larger difference in IAF between the two tasks was associated with a more pronounced task-related change in both TBW and sensitivity. This indicates that some of the differences in TBW and sensitivity between the two tasks are probably related to the shift in alpha frequency during task performance. This probably reflects a narrowing of the TBW and speeding up of alpha in the simultaneity task, which maximizes the demands on the temporal segregation of the visual and tactile signals, compared to the body ownership task that is not a ‘temporal task’ but a task about the illusory feeling of the rubber hand as one’s own or not. However, when we calculated FOOOF-IAF, we did not find any significant differences in the alpha peak between the two tasks. Therefore, we cannot exclude the possibility that the modulation of the alpha peak by the perceptual task demand may be driven by the aperiodic component of alpha frequencies. Thus, our findings about task-related modulations of IAF should be interpreted with caution. (see FOOOF Results).

**Experiment 2: no correlation between alpha power and sensory uncertainty**

We found no significant relationship between sensory uncertainty and alpha power measured during the body ownership judgments both in the left (Pearson’s r = .011; p = .940, Spearman’ ρ: -.031; p = .837) and right parietal lobes (Pearson’s r = .020; p = .895; Spearman’s ρ: -.024; p = .877). Similarly, there was no correlation between alpha power measured during the simultaneity judgments and sensory uncertainty (σ) in the left (Pearson’s r = -.010; p = .947; Spearman’s ρ = -115; p = .446) and right parietal cortex (Pearson’s r = -.023 p = .878; Spearman’s ρ = -117; p = .441) Resting-state alpha power also did not correlate with sensory uncertainty (left: Pearson’s r = -.119; p = .430; Spearman’s ρ = -.055; p = .717; right: Pearson’s r = -.045; p = .768; Spearman’s ρ = -.058; p = .704).

**Experiment 2: no correlation between IAF and prior probability of a common cause between vision and touch**

The prior probability for a common cause between visual and tactile stimuli in the body ownership task did not show any correlation with neither task-related IAF (left parietal lobe: Pearson r = .031; p = .851; Spearman’s ρ = -.033; p = .845; right parietal lobe: Pearson r = .055; p = .742; Spearman’s ρ = -.008; p = .960) and resting-state IAF (left parietal lobe: Pearson r = .187; p = .262; Spearman’s ρ = -.167; p = .315; right parietal lobe: Pearson r = .221; p = .183; Spearman’s ρ = .195; p = .240). Similarly, the prior probability for a common cause in the visuotatcile simultaneity judgments did not show any correlation with neither task-related IAF (left parietal lobe: Pearson r = -.043, p = .797; Spearman’s ρ = .-.052, p = .756; right parietal lobe: Pearson r = -.083; p = .618; Spearman’s ρ = -.105, p = .529) and resting-state IAF (left parietal lobe: Pearson r = .240; p = .146; Spearman’s ρ = .307, p = .060; right parietal lobe: Pearson r = .180; p = .280; Spearman’s ρ = .204; p = .218).

**Supplementary results**

**Experiment 2: FOOOF analysis**

The local maximum power method used in the main manuscript to calculate the IAF has some limitations. First, not all the participants might show a clear peak in the alpha band. Indeed, during task engagement alpha activity is suppressed, and the identification of a frequency band’s peak becomes more challenging and less reliable. Secondly, EEG signal reflects not only the periodic (i.e., oscillatory) component, but also additional background aperiodic activity. This aperiodic activity is present across all frequencies and follows a 1/f power distribution, whereby spectral power decreases as frequency increases^3-6^. Recent works highlight the risk that correlations with the aperiodic component could be erroneously interpreted as relationships with oscillations, especially when focusing on a narrow band of interest^4,6^. To account for these limitations, we implemented the recent "Fitting Oscillations and One Over F" (FOOOF) algorithm to increase the precision of periodic estimates by modeling brain oscillations after excluding the aperiodic component^3^. To address the limitations of the local maximum method in estimating the individual alpha peak, we also implemented the FOOOF algorithm. This allowed us to increase the precision of the oscillation estimates by excluding the aperiodic component of the EEG spectrum. We applied the FOOOF algorithm^3^ to Welch’s power spectral density (PSD) data using the FOOOF/Specparam function in Brainstorm^7^ for MATLAB. This algorithm was implemented separately on each electrode for the alpha frequency range (8–13 Hz) with a sliding time window with 50% overlap to fit spectral peaks, using default parameters: a Gaussian distribution, a minimum number of peaks of 3, a minimum peak height of 2 dB, and a proximity threshold of two standard deviations from the largest peak. To assess the FOOOF model’s performance, we quantified the goodness of fit (R²) for the body ownership, simultaneity judgment, and resting-state conditions across all electrodes of interest. We then extracted peak parameters from the periodic component and calculated the IAF.

**Experiment 2: FOOOF results**

In the resting state condition, FOOOF algorithm successfully identified an alpha peak (8-13 Hz) in most of the 12 electrodes of our ROIs (mean electrodes with a peak: 11,282; range: 0-12; mean fitted peak R^2^ = 0,968; SD = 0,015). In the body ownership judgments, the FOOOF algorithm found a peak on 9,369 electrodes on average (range: 0-12; mean fitted peak R^2^ = 0,973; SD = 0,025) while in the simultaneity judgments the algorithm identified a peak in 9,500 electrodes (range: 0-12; mean fitted peak R^2^ = 0,976; SD = 0,016). Alpha oscillations might be suppressed during perceptual task and peak identification become more challenging, this is probably why we found a greater number of fitted peaks in the resting state as compared to body ownership and simultaneity judgments. Since the FOOOF algorithm identified more peaks in the resting state compared to the perceptual tasks, we conducted two analyses: one focusing on the resting state IAF, including all participants who had at least one alpha peak identified in each ROI (N = 43; mean electrodes with a peak: 11,906 range: 9-12); and another focusing on task-related IAF, which included only participants with at least one alpha peak identified in each ROI for both the body ownership and simultaneity judgment tasks (N = 35; mean electrodes with a peak: 11,319; range: 4-12)

**Experiment 2: correlation between FOOOF-IAF and TBWs**

We observed a negative correlation between parietal resting state FOOOF-IAF and body ownership TBW in both the left (Pearson’s r = -.459; p = .002; Spearman’s ρ = -.561; p < .001 N = 43) and right (Pearson’s r = -.469; p = .002; Spearman’s ρ = -.512; p < .001; N = 43) lobules. Similarly, resting state FOOOF-IAF correlated with simultaneity TBW in the left (Pearson r = -.598; p < .001; Spearman’s ρ = -.656; p < .001; N = 43) and right hemispheres (Pearson’s r = -.540; p < .001, Spearman’s ρ = -.576; p < .001; N = 43). We also found significant correlations when considering task-related FOOOF-IAF in participants whose alpha peak was successfully fitted during the perceptual judgments. Specifically, IAF correlated with body ownership TBW in both left (Pearson’s r = -.556; p =.001, Spearman’s ρ = -.551; p = .001; N = 35; **Fig S4a**) and right parietal cortex (Pearson’s r = -.569; p < .001, Spearman’s ρ = -.562 < .001; N = 35). Similarly, task related IAF correlated with simultaneity TBW in both left (Pearson’s r = -.646; p < .001, Spearman’s ρ = -.617 < .001; N = 35; **Fig S4c**) and right hemispheres (Pearson’s r = -.648; p < .001, Spearman’s ρ = -.631; p < .001; N = 35)

**Experiment 2: correlation between FOOOF-IAF and sensitivity**

We found a positive and significant correlation between resting-state FOOOF-IAF and sensitivity to body ownership in both hemisphere (left: Pearson’s r = .405, p = .007; Spearman’s r = .439, p = .003; right: Pearson’s r: .426, p = .004; Spearman’s ρ = .435, p = .004; N = 43). Resting state IAF correlated also with sensitivity to visuotactile simultaneity in both left (Pearson’s r = .589, p < .001; Spearman’s ρ = .638, p < .001; N = 43) and right hemisphere (Pearson’s r = .569, p < .001; Spearman’s ρ = .598, p < .001; N = 43). We also found positive correlations when considering task-related FOOOF-IAF. Specifically, task related IAF correlated with body ownership sensitivity in both left (Pearson’s r = .614, p < .001; Spearman’s ρ = .581, p < .001; N = 35; **Fig S4b**) and right hemispheres (Pearson’s r = .590, p < .001; Spearman’s ρ = .565, p < .001; N = 35). Task related FOOOF-IAF also significantly correlated with visuotactile simultaneity sensitivity in both left (Pearson’s r = .715, p < .001; Spearman’s ρ = .664, p < .001; N = 35; **Fig S4d**) and right parietal lobules (Pearson’s r = .644, p < .001; Spearman’s ρ = .629, p < .001; N = 35)

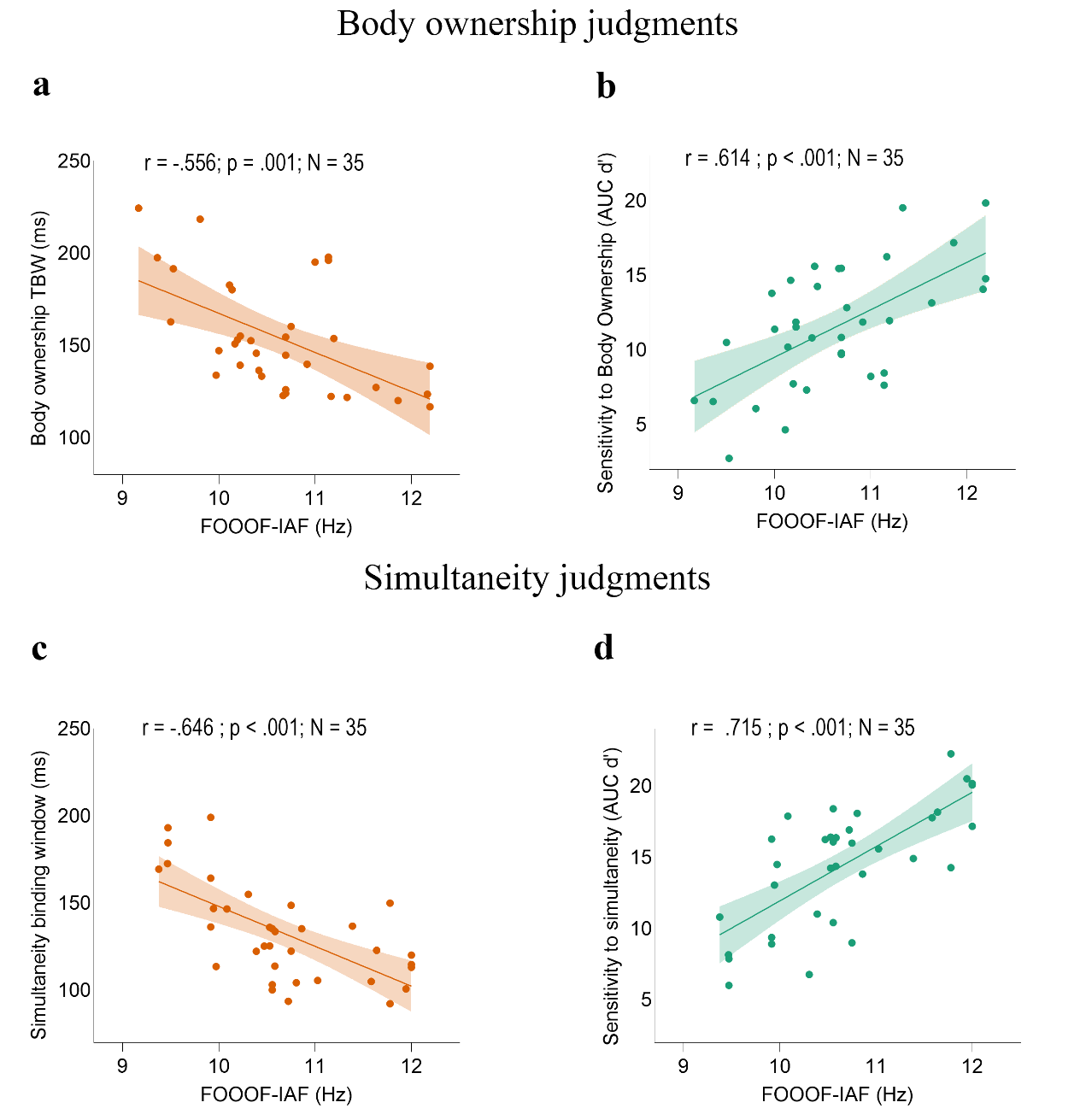

**Fig S4 | Correlations between task-related FOOOF-Individual Alpha Frequency in the left parietal lobe and perceptual performance in Experiment 2. a, c,** Correlation between task-related FOOOF-IAF measured in the left posterior parietal cortex and body ownership and visuotactile simultaneity TBWs. **b, d,** Correlation between task-related FOOOF-IAF in the left posterior parietal cortex and sensitivity to body ownership and to visuotactile simultaneity (**d**). The solid line represents the best-fitting regression. The shaded region reflects the 95%

**Experiment 2: correlation between FOOOF-IAF and sensory uncertainty**

When considering the relationship between resting state FOOOF-IAF and sensory uncertainty we found a significant correlation in the left hemisphere (Pearson’s r = -.319, p = .037; Spearman’ ρ = -.416; p = .006; N = 43) and a close to significant correlation in the right hemisphere (Pearson’s r = -.299, p = .051; Spearman’ ρ = -.352; p = .021; N = 43). Task evoked FOOOF-IAF measured during body ownership judgments, significantly correlated with the level of sensory uncertainty in both left (Pearson’s r = -.549, p = .001; Spearman’ ρ = -.571, p < .001; N = 35; **Fig S5a**) and right (Pearson’s r = -.471, p = .004; Spearman’ ρ = -.482, p = .003; N = 35) parietal lobes. Similarly, FOOOF-IAF measured during simultaneity judgments correlated with the level of sensory uncertainty in both left (Pearson’s r = -.541, p = .001; Spearman’ ρ = -.526, p = .001; N = 35; **Fig. S5b**) and right (Pearson’s r = -.537, p = .001; Spearman’ ρ = -.492, p = .003; N = 35) hemisphere.

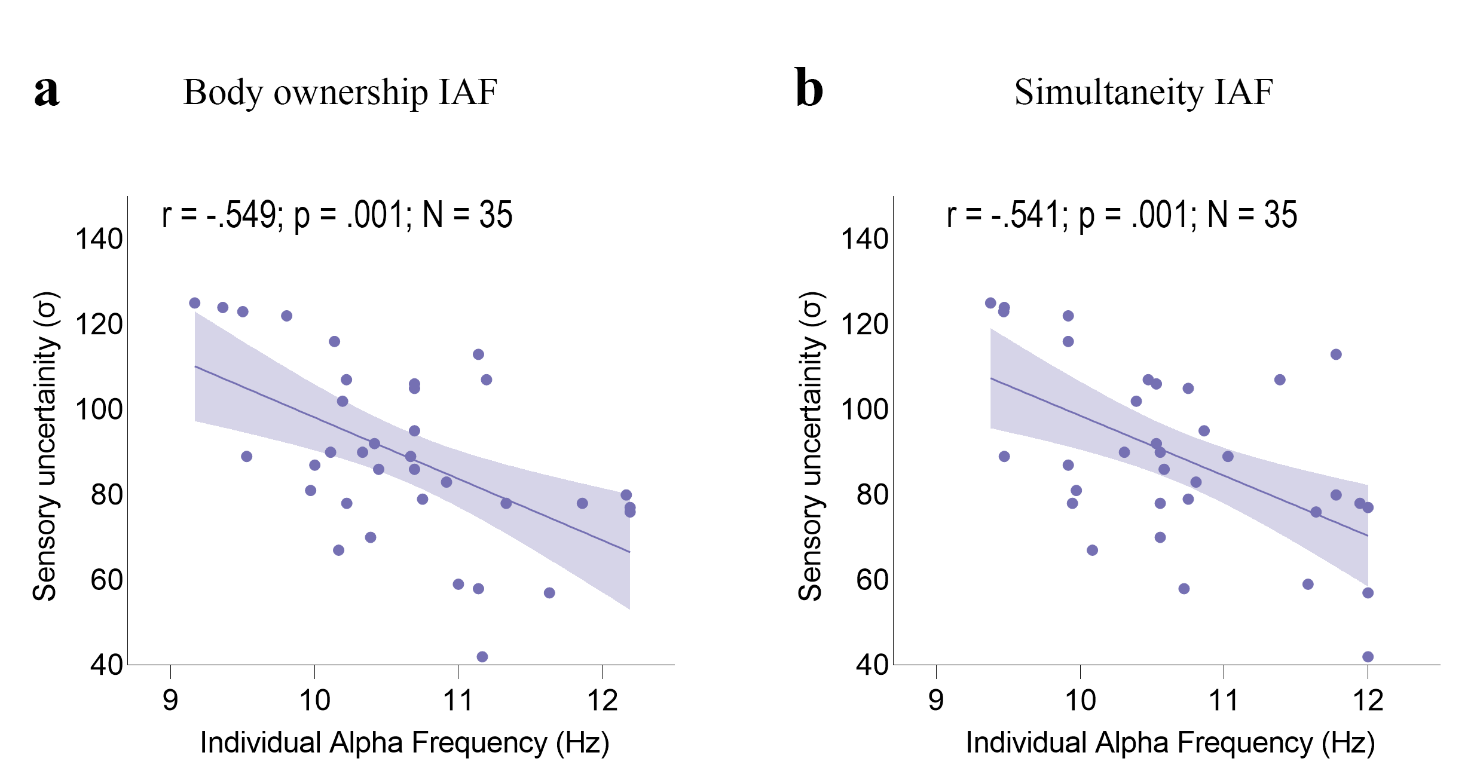

**Fig. S5 | Correlations between task-related FOOOF-Individual Alpha Frequency (IAF) in the left parietal lobe and the level of sensory uncertainty (σ) in Experiment 2. a, b,** Correlation between sensory uncertainty (σ) and task-related IAF measured during body ownership and simultaneity judgments. The solid line represents the best-fitting regression. The shaded region reflects the 95% confidence interval

**Experiment 2: IAF modulation between body ownership and simultaneity judgments.**

When computing the IAF using the local maximum method, we found that IAF measured during body ownership judgments was slower as compared to the IAF measured during simultaneity judgments. However, when controlling for the aperiodic component we found no significant differences in the FOOOF-IAF between body ownership and simultaneity judgments (F_1,34_ = 0.671, p = .418, η_p_² = .019). Moreover, we found no significant differences between left and right parietal lobes (F_1,34_ = 3.398, p = .074, η_p_² = .091).

**Experiment 3: tACS effect in body ownership correlates with tACS effect in simultaneity judgements**

Because alpha tACS modulated both body ownership and simultaneity judgments, we examined whether participants whose simultaneity judgments were more affected by tACS were also those whose body ownership judgments were most influenced by the brain stimulation. To test this, we subtracted the TBW in the low alpha tACS condition from the TBW in the sham tACS in both ownership and simultaneity judgments to obtain a measure of each participant's low tACS effect. The same procedure was repeated for the high alpha tACS condition, subtracting the TBW obtained in the sham from the TBW obtained in the high alpha tACS condition in both body ownership and simultaneity judgments. Thus, greater differences indicated larger alpha tACS effects in each condition (low and high). We found positive correlations between the tACS effects in the body ownership TBW and the simultaneity TBW, respectively. That is, participants whose body ownership judgments were more affected by tACS also had their simultaneity judgments that were more influenced by the brain stimulation. This was true for both low alpha-tACS (Pearson’s r = .728, p <.001; Spearman’s ρ = .702, p < .001; **Fig. S6 a**), and high alpha-tACS conditions (Pearson’s r = .547, p = .002; Spearman’s ρ = .545, p = .002; **Fig. S6 b**).

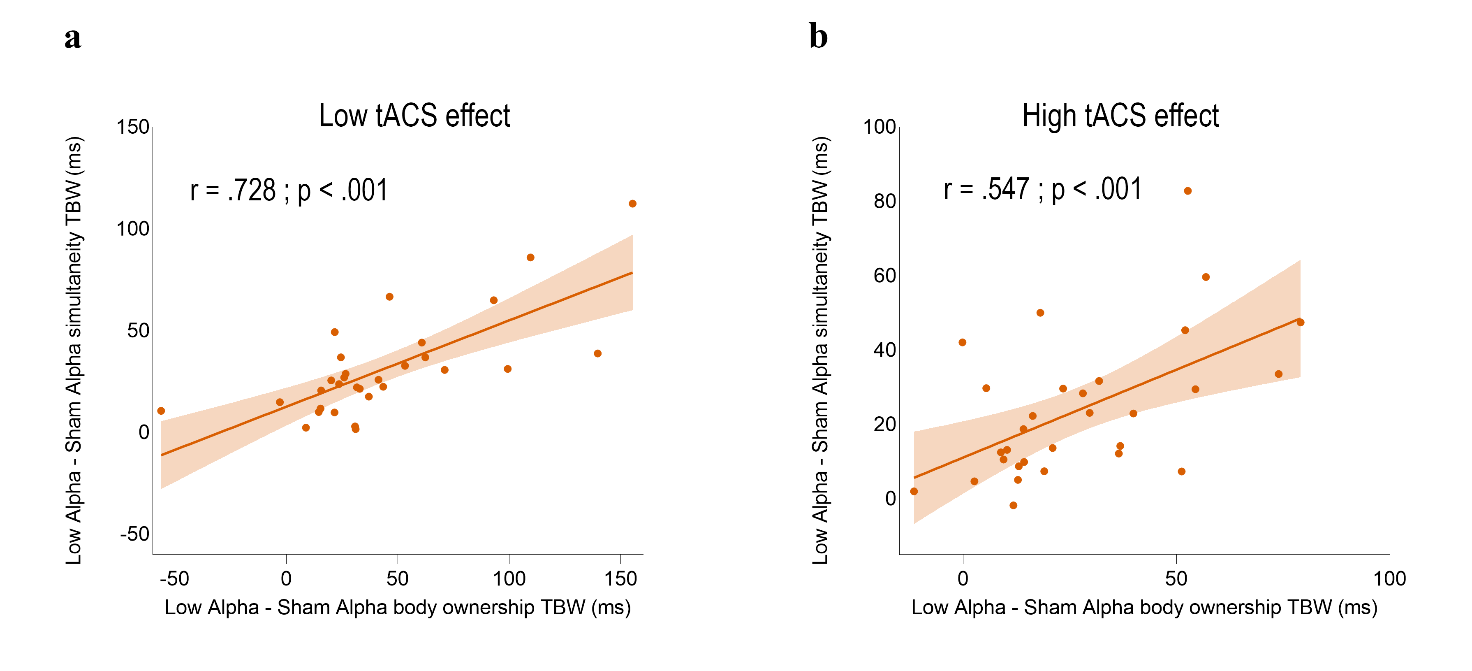

**Fig. S6 | correlation between tACS effects in body ownership and simultaneity judgments a,** Correlation between low alpha tACS effect in body ownership and low alpha tACS effect in visuotactile simultaneity. **b**, Correlation between high alpha tACS effect in body ownership and high alpha tACS effect in visuotactile simultaneity. The shaded region reflects the 95% confidence interval. AUC denote the area under the curve of d’ scores measured for each asynchrony.

**Experiment 3: no tACS effect on bias**

In contrast, there was no significant tACS effect on bias in either body ownership (F_2,58_ = 0.528; p = .593; η^2^_p_ = 0.018) or simultaneity judgments (F_2,58_ = 0.945; p = 0.395; η^2^_p_ = 0.032). The bias changed only as a function of the asynchrony, as participants tended to be more liberal to report body ownership (F_2, 58_ = 352.733; p < .001; η^2^_p_ = 0.924) or the simultaneity (F_6,174_ = 407,642; p < .001; η^2^_p_ = 0.934) at smaller asynchronies.

**Experiment 3: correlation between body ownership and simultaneity TBWs and their perceptual sensitivities**

Experiment 3 successfully replicated the correlations between body ownership and simultaneity TBWs found in the previous experiments. Body ownership TBW correlated with simultaneity TBW in low alpha (Pearson’s r = .845; p < .001; Spearman’s ρ = .752; p < .001; **Fig S7a**), high alpha tACS conditions (Pearson’s r = .670; p < .001; Spearman’s ρ = .609; p < .001; **Fig S7c**) and sham tACS condition (Pearson’s r = .748; p < .001; Spearman’s ρ = .781; p < .001; **Fig S7e**). Moreover, body ownership sensitivity correlated with simultaneity sensitivity in low alpha (Pearson’s r = .525; p = .003; Spearman’s ρ = .494; p = .006; **Fig S7b**), high alpha (Pearson’s r = .556; p = .001; Spearman’s ρ = .552; p = .002; **Fig S6d**) and sham condition (Pearson’s r = .681; p < .001; Spearman’s ρ = .625; p < .001; **Fig. S7f**). Similarly to previous experiment, paired t-tests revealed that body ownership TBW was wider than simultaneity TBW in sham (t = 6.206; p < .001; d = 1.133), low alpha (t = 6.805; p < .001; d = 1.242) and high alpha (t = 5.683; p < .001; d = 1.037) tACS conditions. The ANOVAs conducted with stimulation types and asynchronies as factors on body ownership and simultaneity sensitivity indices revealed that, beyond the main effect of stimulation highlighted in the manuscript, both body ownership sensitivity (F_2, 58_ = 353.612, p < .001, η²ₚ = .924) and simultaneity sensitivity (F_2, 58_ = 407.642, p < .001, η²ₚ = .934) significantly increased with larger asynchronies.

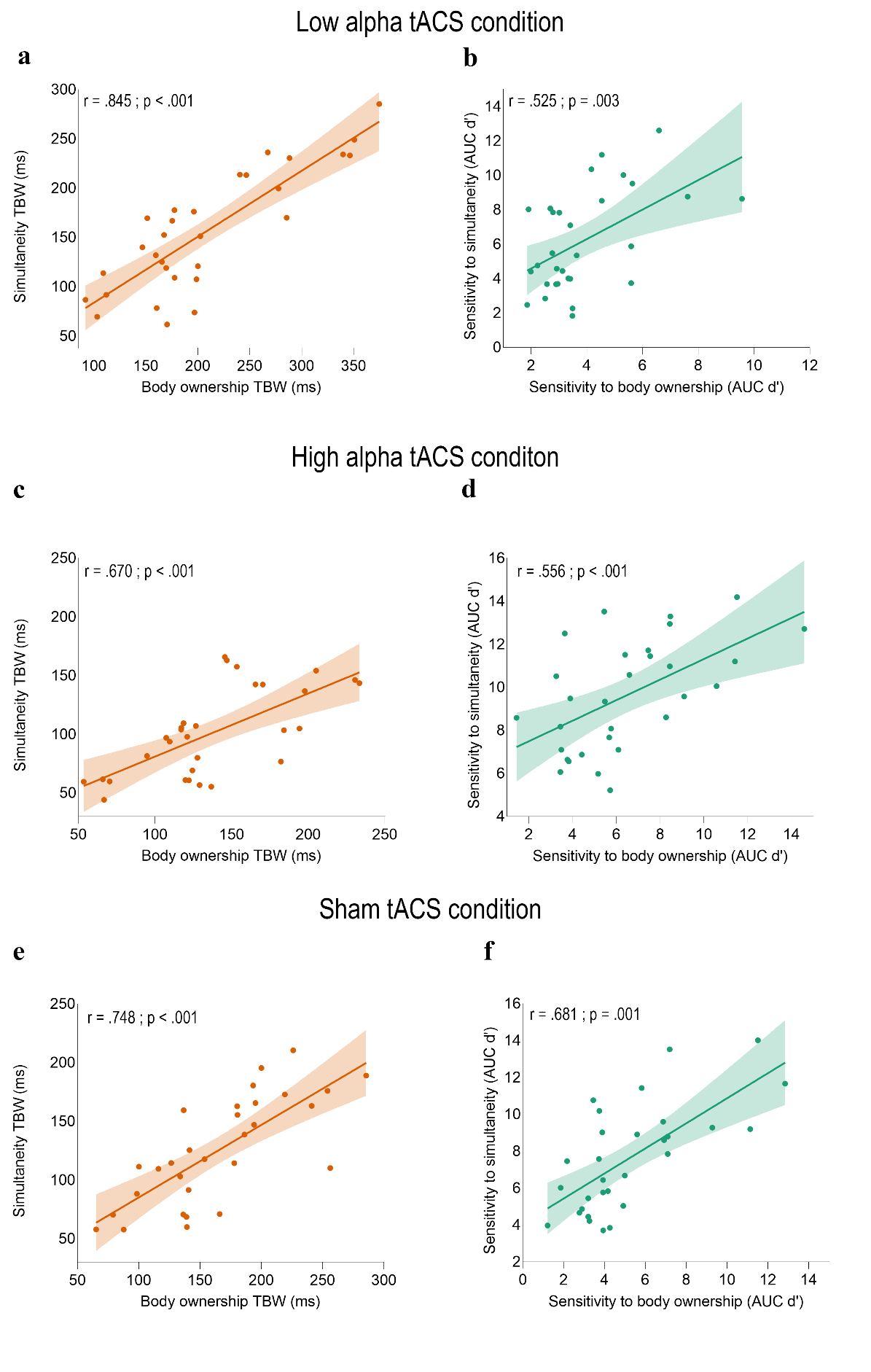

**Fig. S7 | Behavioural correlations in Experiment 3 a, c, e,** Correlations between body ownership and simultaneity judgements’ TBWs as measured in the low alpha (**a**), high alpha (**d**) and sham (**e**) tACS conditions. **b, d, f,** Correlations between sensitivity to body ownership and simultaneity in low alpha (**b**), high alpha (**c**) and sham (**f**) tACS conditions

**Experiment 3: survey of sensations related to transcranial Alternating Current Stimulation**

To investigate the effectiveness of participants’ blindness relative to the brain stimulation conditions, we tested for any difference in participants’ self-reported physical sensations between tACS conditions and sham. To this aim, participants completed a validated questionnaire^8^ assessing tACS-related sensations at the end of each experimental session (**Attachment 2**). An initial question evaluated if participants believed that they had received real or sham stimulation. The next questions assessed seven specific physical sensations usually associated with electrical brain stimulation (i.e., itchiness, pain, burning, warmth, iron taste and fatigue). The final question examined participants’ view that the sensations related to tACS may have changed their performance in the tasks.

Each answer was scored on a value ranging from 1 (‘I definitely did not receive real brain stimulation / I felt no sensations / performance did not change at all’) to 5 (‘I definitely received real brain stimulation / I strongly felt one or more sensations / performance changed a lot’). To assess if participants believed if they received real or sham stimulations, we compared the score of the first question with a one-way ANOVA with brain stimulation (low alpha, high alpha and sham tACS) as main factor. The ANOVA showed no main effect of brain stimulation (F_2,58_ = .585; p = 0.585; η_p_^2^ = 0.018), indicating that participants could not reliably distinguish between real or sham stimulation conditions.

Next, differences in perceived physical sensations across tACS conditions were examined in a repeated measure ANOVA with brain stimulation (low alpha, high alpha, sham-tACS) and physical sensations (itchiness, pain, burning, warmth, pinching, iron taste and fatigue) as factors. The analysis revealed a main effect of physical sensations (F_6, 174_ = 8.304; p < 0.001; η_p_^2^ = 0.223) because participants reported in general significantly stronger itchiness as compared to other sensations, i.e., pain, burning and iron taste (p < 0.010 for all comparisons) and stronger fatigue as compared to pain, burning, warmth, iron taste (p < 0.003 for all comparisons). However, neither the stimulation condition (F_2,58_ = 0.151; p = 0.860; η_p_^2^ = 0.005) neither its interaction with physical sensations were significant (F_12, 348_ = 0.928; p = 0.518; η_p_^2^ = 0.031), indicating that there were no differences in the perceived physical sensations between the three tACS conditions. Moreover, no participants reported other physical sensations, such as phosphenes or flickering sensations. Finally, participants’ view on whether performance may have been changed following stimulation did not differ across conditions (one-way ANOVA, F_2,58_ = 0.473; p = 0.625; η_p_^2^ = 0.016). Overall, these results clearly indicate that participants could not distinguish between real and sham stimulation conditions. Specifically, real stimulation did not differ from sham in terms of reported physical sensations and participants’ perceived change in performance.

**Attachment 1**

| Q1 | It seemed as if I were feeling the touch in the location where I saw the rubber hand touched |
| --- | --- |
| Q2 | It seemed as though the touch I felt was caused by the stick touching the rubber hand |
| Q3 | I felt as if the rubber hand were my hand |
| Q4 | It felt as if my (real) hand were drifting up (towards the rubber hand) |
| Q5 | It seemed as if I might have more than one right hand or arm |
| Q6 | It seemed as if the touch I was feeling came from somewhere between my own hand and the rubber hand |
| Q7 | It felt as if my (real) hand was turning ‘rubbery’. |
| Q8 | It appeared (visually) as if the rubber hand were drifting towards my hand. |
| Q9 | The rubber hand began to resemble my own (real) hand, in terms of shape, skin tone, freckles or some other visual feature. |

**Attachment 2**

**Survey of sensations related to transcranial Alternate Current Stimulation**

In your opinion, have you received real brain stimulation during today’s experimental session?

Please answer accordingly to the following scale

- **1= I DEFINITELY received real brain stimulation**
- **2= I PROBABLY received real brain stimulation**
- **3= I’m NOT ABLE TO DETERMINE if I have received real brain stimulation**
- **4= I PROBABLY did NOT receive real brain stimulation**
- **5= I DEFINITELY did NOT receive real brain stimulation**

**□ 1	□2	□3	□4	□5**

Have you experienced any discomfort or annoyance during the stimulation?

Please answer to the following questions regarding the different sensations, indicating the degree of intensity of your discomfort according to the following scale:

- **None = I have not felt the described sensations**
- **Mild = I have mildly felt the described sensations**
- **Moderate= I have felt the described sensations**
- **Considerable= I have felt the described sensations to a considerable degree**
- **Strong= I have strongly felt the described sensations**

Itchiness: □ None	□ Mild	□Moderate	□Considerable	□Strong

Pain: □ None	□ Mild	□Moderate	□Considerable	□Strong

Burning: □ None	□ Mild	□Moderate	□Considerable	□Strong

Warmth/Heat: □ None	□ Mild	□Moderate	□Considerable	□Strong

Pinching: □ None	□ Mild	□Moderate	□Considerable	□Strong

Iron taste: □ None	□ Mild	□Moderate	□Considerable	□Strong

Fatigue: □ None	□ Mild	□Moderate	□Considerable	□Strong

Other:_____________: □ None	□ Mild	□Moderate	□Considerable	□Strong

When did the discomfort begin?

□ At the beginning of the experiment □About the middle of the experiment □Towards the end of the experiment

How long did it last?

□It stopped soon □It stopped in the middle of the block □It stopped at the end of the block

How much these sensations affect your performance in the tas

□ Not at all	□ A little	□ Considerably	□ Much	□ Very much
